# GENOME ASSEMBLIES FOR 28 PHAEOSPHAERIACEAE ISOLATES FROM CANADA

**DOI:** 10.64898/2026.11.02.763635

**Authors:** Janice Bamforth, Mohamed Abdel-Fattah, Ryan Gourlie, Reem Aboukhaddour, Sean Walkowiak

## Abstract

Glume blotch and Septoria Tritici Blotch (STB) are destructive crop diseases caused by fungal pathogens formerly classified under the genus *Septoria*. Recent taxonomical revisions have led to the reclassification and renaming of these fungi, some of which are now under the genus *Parastagonospora*, family Phaeosphaeriaceae, order Pleosporales, and class Dothideomycetes. In this study, we performed whole genome sequencing and assembly for 28 Phaeosphaeriaceae isolates from Canada. Sequencing was performed using a combination of Oxford Nanopore Technologies (ONT) long-read and Illumina short-read sequencing platforms. Most of the genomes were very complete, BUSCO >99%, with the genomes assemblies derived from ONT sequencing having very few contigs that likely encompass entire chromosomes or chromosome arms. To determine the relationship between these isolates and similar fungi that have been previously characterized, we performed whole genome alignments to 2,668 other Dothidiomycete genomes obtained from NCBI using NUCmer. Comparative genomic analysis revealed that 27 of the isolates clustered discretely with either *Parastagonospora nodorum, or Parastagonospora pseudonodorum (Parastagonospora avenae f. sp. tritici*), with one genome having no other similar genome in NCBI which we hypothesize to be the first genome assembly of *Phaeosphaeria pontiformis*. Both *P. nodorum* and *P. pseudonodorum* species were isolated from wheat and barley and only the latter was isolated from durum, while the *P. pontiformis* was isolated from wheat. In summary, we sequenced and assembled the genomes of 28 isolates of Phaeosphaeriaceae spp. from Canada and the genomes add to our foundational knowledge of these plant pathogens and efforts to better understand their diversity.

## Introduction

Cereals, including wheat and barley, are important staple foods that contribute significantly to human caloric and protein intake. Yield and quality losses of cereals are of significant food security concern, particularly in the context of the growing human population and increased pressures from abiotic and biotic stress brought on by climate change (Robles-Zazueta et al., 2024). Phaeosphaeriaceae spp. are important fungal pathogens of cereals worldwide that can cause foliar diseases that result in yield losses and can contaminate and reduce the quality of grain (Ficke et al., 2018). Monitoring of plant pathogen populations is an important component of disease management programs and allows pathologists and breeders to better understand the dynamics of diseases and emergence of new pathogen strains.

While traditional disease monitoring and pathogen identification have relied on culture, tissue, and cell morphology, modern DNA-based methods are creating universal standards on which to identify and classify microbes. DNA barcodes can be used to classify species as well as subdivide the species into different populations and lineages (Ueng et al., 1992; Ueng & Chen, 1994; Ueng et al., 1995). However, DNA barcodes are not universally applicable to all fungi and do not capture genes that may be contributing to disease (Thines et al., 2018). Studies have identified a number of different virulence factors within Phaeosphaeriaceae spp, including ToxA, Tox1, and Tox3 that contribute to disease and plant-pathogen interactions, monitoring of which has enhanced our understanding of the distribution of disease related genes (Hafez et al., 2020). Specialized genomic elements, starships, within the fungi have been identified to facilitate horizontal gene transfer of some of disease genes across fungi, including the ToxA virulence factor, supporting the need for whole genome information to understand the disease potential and risk of different fungal isolates (Griem-Krey et al., 2026). Whole genome sequencing is perhaps the most thorough method classifying organisms and assessing their diversity and has been helpful in understanding pathogen emergence and identification of new taxa (Croll et al., 2021). Here, we sought to build upon the foundation of genomic resources by performing whole genome sequencing of isolates of Phaeosphaeriaceae from Canada, which will add to our collective knowledge of these pathogens and their diversity.

## Methods

Two panels of fungal isolates were used in this study. The first panel was sequenced using both Illumina and long-read Oxford Nanopore Technologies (ONT) platforms, while the second panel was sequenced using only short-read Illumina technologies.

For the first panel, ten fungal isolates from the years 2005 to 2015 were selected for whole genome sequencing from the culture collection of the Canadian Grain Commission, with most isolates being isolated from damaged grain or ergot bodies contaminated with other fungi (Table 1). One isolate was from the Canadian Collection of Fungal Cultures (DAOMC) culture collection, DAOMC 182701, and overlapped with the second panel. Cultures were recovered and grown on potato dextrose agar at room temperature for approximately five days. Mycelia were harvested and frozen at −140 °C in an ultralow temperature freezer. Mycelia were ground in a mortar and pestle, and DNA was extracted and precipitated using cetyltrimethylammonium bromide (CTAB) and chloroform-isoamyl alcohol methods. Purification and size selection using the BluePippin High Pass Plus kit was used for samples with low quality, according to manufacturer protocols. Whole genome sequencing was performed on an Oxford Nanopore Technologies (ONT) GridIon instrument using R9.4.1 flow cells and the Rapid Sequencing Kit (SQK-RAD004). In parallel, samples were sequenced using Illumina iSeq 150 bp paired-end technologies, using manufacturer recommended protocols. Genome assemblies were generated from the ONT sequencing reads using Flye (Kolmogorov et al., 2019) and polished with the Illumina sequencing reads using BWA-MEM (Li, 2013) and Pilon (Walker et al., 2014).

**Table 1.** List of fungal isolates.

| **Isolate Name** | **Year** | **Location** | **Source** | **Species** |
|---|---|---|---|---|
| Alt01_033 | 2013 | Unknown | Winter Wheat | *P. nodorum* |
| DAOMC 182701 | 1981<sup>a</sup> | Prince Edward Island | Spring Wheat | *P. nodorum* |
| Fus15_20 | 2013 | Prince Edward Island | Barley | *P. nodorum* |
| MS02_31 | 2005 | Alberta | Spring Wheat | *P. nodorum* |
| MS02_32 | 2005 | Alberta | Unknown | *P. nodorum* |
| Alt01_021 | 2013 | Unknown | Barley | *P. pseudonodorum* |
| Fus15_19 | 2012 | Saskatchewan | Durum Wheat | *P. pseudonodorum* |
| Fus19_41 | 2015 | Alberta | Durum Wheat (ergot sclerotia) | *P. pseudonodorum* |
| Fus19_42 | 2015 | Manitoba | Spring Wheat (ergot sclerotia) | *P. pseudonodorum* |
| Fus19_43 | 2015 | Manitoba | Spring Wheat (ergot sclerotia) | *P. pseudonodorum* |
| G211-5<sup>b</sup> | 2018 | Alberta | Spring Wheat | *P. nodorum* |
| S419-22F<sup>b</sup> | 2019 | Saskatchewan | Spring Wheat | *P. nodorum* |
| 06-SN-004<sup>b</sup> | 2006 | Manitoba | Spring Wheat | *P. nodorum* |
| B313-13<sup>b</sup> | 2018 | Alberta | Spring Wheat | *P. nodorum* |
| C316-42F<sup>b</sup> | 2020 | Alberta | Spring Wheat | *P. nodorum* |
| DAOMC 162797<sup>b</sup> | 1977 | Ontario | Spring Wheat | *P. nodorum* |
| BC-07<sup>b</sup> | 2022 | British Columbia | Spring Wheat | *P. nodorum* |
| DAOMC 182701<sup>b</sup> | 1981 | Prince Edward Island | Spring Wheat | *P. nodorum* |
| S415-3<sup>b</sup> | 2018 | Saskatchewan | Spring Wheat | *P. nodorum* |
| C113-75F<sup>b</sup> | 2021 | Alberta | Spring Wheat | *P. nodorum* |
| C106-15<sup>b</sup> | 2018 | Alberta | Spring Wheat | *P. pseudonodorum* |
| 06-SN-027<sup>b</sup> | 2006 | Manitoba | Spring Wheat | *P. pseudonodorum* |
| B301-14<sup>b</sup> | 2018 | Alberta | Spring Wheat | *P. pseudonodorum* |
| SK71-10<sup>b</sup> | 2022 | Saskatchewan | Spring Wheat | *P. pseudonodorum* |
| 07-SN-019<sup>b</sup> | 2007 | Manitoba | Spring Wheat | *P. pseudonodorum* |
| 07-SN-012<sup>b</sup> | 2007 | Manitoba | Spring Wheat | *P. pseudonodorum* |
| 07-SN-017<sup>b</sup> | 2007 | Manitoba | Spring Wheat | *P. pseudonodorum* |
| B102-44bF<sup>b</sup> | 2019 | Alberta | Spring Wheat | Pleosporales sp. |

<sup>a</sup> DAOMC 182701 was first deposited in the DAOMC collection of Agriculture and Agri-Food Canada in 1981, the isolation date is unknown – this isolate is part of both panels
<sup>b</sup> indicates the second panel of isolates (Illumina sequencing only)

For the second panel, fungal isolates were recovered from node and grain samples collected between 2006 and 2022, along with reference isolates obtained from DAOMC, as indicated in Table 1. The preparation fungal tissue for DNA extraction first involved culturing on V8 potato dextrose agar medium. Cultures were incubated under continuous light at room temperature (∼25 °C) for at least 5 days. Next, approximately 5 mL of autoclaved distilled water was added to the plate, and the mycelium is scraped with a loop or crushed using the back of a glass tube. The scraped mycelial suspension was then transferred to a fresh V8-PDA plate and spread evenly using an L-shaped spatula. Both the old and new plates were subsequently maintained under continuous light at room temperature (∼25 °C) without parafilm. After 3 to 4 days, dark spots begin to appear on both plates. One to two days after the appearance of these black spots, 5 to 10 mL of autoclaved distilled water was added to the plates, and the surfaces were scraped to collect spores. The collected spores and mycelium were then transferred into 250 mL flasks containing 100 mL of sterile dilute potato dextrose broth, at one quarter concentration, or low Fries medium, and grown in darkness at room temperature. The top mycelial mat formed in the liquid medium was carefully removed and washed twice with sterile distilled water to eliminate excess growth medium. Finally, the collected mats were freeze-dried and stored at −20 °C for subsequent DNA extraction. For short-read sequencing, genomic DNA was extracted as described in Gourlie et al. (2022) using the ‘Genomic-tip 100/G’ kit (Qiagen, Mississauga, ON, Canada) and sequenced with 150 bp paired-end, 400 bp inserts at ∼100X□coverage with Illumina HiSeq (Gourlie et al., 2022). Shovill with SPAdes was used for de novo assembly (Seemann, 2026).

For the assemblies from both panels, assembly statistics were generated using QUAST and BUSCO (database: pleosporales_odb12). Genes were predicted using Augustus v3.5.0 using the species parameters for *Fusarium*. To determine the relationship between these isolates and similar fungi that have been previously characterized, we aligned the newly assembled genomes to 2,668 Dothidiomycete genomes obtained from NCBI using MUMmer, using the default parameters of NUCmer. Alignments were then filtered and summary statistics generated using dnadiff. The ITS region of B102-44bF was also extracted and used in BLASTn analysis to facilitate a putative identification.

## Results and Conclusions

Agriculture and Agri-Food Canada (DAOMC) and the Canadian Grain Commission have large fungal culture collections of historically important fungi including the Phaeosphaeriaceae spp. sequenced here. Genome assembly of the isolates from the first panel that used a combination of ONT long-reads and Illumina short-reads resulted in assemblies with 22 to 48 contigs and an N50 of 13.8 to 18.1 Mbp, with most of the assembly represented in 8 to 11 contigs (Table 2).

**Table 2.** Genome Assembly Statistics.

| Assembly | Contigs | Length (Mbp) | GC (%) | N50 | N90 | L50 | L90 | BUSCO (%) | Augustus Predicted Genes |
|---|---|---|---|---|---|---|---|---|---|
| Alt01_033 | 30 | 36.6 | 50.45 | 1733257 | 1078487 | 8 | 20 | 99.1 | 11931 |
| DAOMC 182701<sup>a</sup> | 48 | 37.1 | 50.28 | 1393068 | 528909 | 9 | 23 | 99.1 | 12024 |
| Fus15_20 | 37 | 38.0 | 50.23 | 1726429 | 1139524 | 9 | 20 | 99.1 | 12196 |
| MS02_31 | 26 | 37.4 | 50.45 | 1520991 | 1125547 | 9 | 20 | 99.2 | 12066 |
| MS02_32 | 35 | 36.4 | 50.52 | 1380244 | 804722 | 11 | 24 | 99.1 | 11923 |
| Alt01_021 | 26 | 37.6 | 51.91 | 1815309 | 1240271 | 8 | 18 | 99.3 | 12572 |
| Fus15_19 | 31 | 37.5 | 51.92 | 1643201 | 1214777 | 8 | 19 | 99.4 | 12603 |
| Fus19_41 | 24 | 37.2 | 51.91 | 1618358 | 1244080 | 8 | 18 | 99.3 | 12452 |
| Fus19_42 | 23 | 37.3 | 51.93 | 1648760 | 1288050 | 8 | 18 | 99.4 | 12573 |
| Fus19_43 | 22 | 37.5 | 51.92 | 1656275 | 1255597 | 8 | 18 | 99.4 | 12580 |
| G211-5<sup>b</sup> | 584 | 37.1 | 50.46 | 663782 | 200012 | 17 | 55 | 99.1 | 12111 |
| S419-22F<sup>b</sup> | 400 | 36.7 | 50.39 | 830204 | 219087 | 15 | 49 | 99.1 | 11942 |
| 06-SN-004<sup>b</sup> | 1017 | 37.2 | 50.22 | 778511 | 188623 | 18 | 53 | 99.1 | 11951 |
| B313-13<sup>b</sup> | 322 | 37.3 | 50.14 | 770179 | 301083 | 18 | 48 | 99.1 | 12002 |
| C316-42F<sup>b</sup> | 5922 | 42.4 | 48.78 | 38556 | 4353 | 280 | 1517 | 97.2 | 13064 |
| DAOMC 162797<sup>b</sup> | 375 | 37.3 | 50.01 | 542555 | 255777 | 20 | 58 | 99.1 | 11914 |
| BC-07<sup>b</sup> | 3094 | 39.4 | 49.82 | 194829 | 17963 | 65 | 292 | 99.2 | 12157 |
| DAOMC 182701<sup>b</sup> | 402 | 37.7 | 50.09 | 671127 | 221207 | 20 | 56 | 99.2 | 12060 |
| S415-3<sup>b</sup> | 415 | 36.7 | 50.4 | 696898 | 219172 | 18 | 56 | 99.1 | 11953 |
| C113-75F<sup>b</sup> | 886 | 36.3 | 50.49 | 788387 | 223527 | 14 | 43 | 99.2 | 11865 |
| C106-15<sup>b</sup> | 1143 | 36.0 | 51.96 | 343987 | 89704 | 33 | 108 | 99.3 | 12214 |
| 06-SN-027<sup>b</sup> | 28497 | 39.3 | 52.07 | 3412 | 1122 | 3261 | 10272 | 67.8 | 16967 |
| B301-14<sup>b</sup> | 1934 | 37.0 | 51.98 | 270983 | 56692 | 35 | 141 | 99.4 | 12636 |
| SK71-10<sup>b</sup> | 1710 | 36.5 | 51.92 | 337735 | 66535 | 34 | 126 | 99.3 | 12399 |
| 07-SN-019<sup>b</sup> | 1276 | 36.2 | 51.95 | 360234 | 76377 | 32 | 107 | 99.3 | 12309 |
| 07-SN-012<sup>b</sup> | 132 | 40.3 | 51.61 | 1541303 | 1084563 | 9 | 21 | 99.3 | 13260 |
| 07-SN-017<sup>b</sup> | 167 | 40.0 | 51.79 | 1650910 | 1107255 | 9 | 20 | 99.3 | 13258 |
| B102-44bF<sup>b</sup> | 1495 | 39.1 | 50 | 712066 | 148106 | 18 | 66 | 99.2 | 12529 |

<sup>a</sup> DAOMC 182701 was first deposited in the DAOMC collection of Agriculture and Agri-Food Canada in 1981, the isolation date is unknown – this isolate is part of both panels
<sup>b</sup> indicates the second panel of isolates (Illumina sequencing only)

These contigs likely represent entire chromosomes or chromosome arms, given that *Parastagonospora nodorum* is reported to have 23 chromosomes (Bertazzoni et al., 2021; McDonald et al., 2019). The second panel, which was sequenced only using Illumina short-reads resulted in assemblies with 132 to 28497 contigs and an N50 of 3412 bp to 1.65 Mbp with most of the assembly being represented in 9 to 3261 contigs (Table 2). The notable outlier in the assemblies was 06-SN-027, which was more fragmented. While more fragmented than the genome assemblies from the first panel, most of the assemblies from the second panel had high N50 values, in the hundreds of kilobases to megabase pair range, indicating that they are of reasonable quality. The total lengths of the assemblies ranged from 36.0 to 42.4 Mbp, with most assemblies being between 36.0 and 38.0 Mbp (Table 2), which is the expected genome size for *Parastagonospora* species (Bertazzoni et al., 2021; McDonald et al., 2019). Analysis of BUSCO genes from Pleosporales indicate that most of the genomes were >99% complete, with the exception of 06-SN-027, likely due to the assembly being more fragmented. Augustus was able to predict between 11,865 and 16,967 genes, with most isolates having between 11,000 and 13,000 predicted genes (Table 2). This is near, but slightly below previously reported gene counts of 13,869 for *Parastagonospora nodorum*; the underestimation may be due to genes being missed by Augustus (Bertazzoni et al., 2021; McDonald et al., 2019). A more thorough gene projection analysis from related species, in combination with RNA and isoform sequencing may be beneficial in the future to fully catalogue the genes for each genome. Nevertheless, the data cumulatively support that the genomes were expected genome lengths for the target species, are assembled into few contigs, particularly for the assemblies from the first panel, and contain a high level of complete BUSCO genes, supporting that they are of high quality and completeness. The notable exception was 06-SN-027, which was more fragmented and had fewer complete BUSCO genes.

Whole genome alignments revealed that most of the genomes had a very high percent genome alignment (>95%) to genomes from public repositories (Supplemental Table 1), with the exception of B102-44bF. Most genomes had high genome alignment to either *P. pseudonodorum (P. avenae f. sp. tritici*) or *P. nodorum* genomes, and had high percent genome alignment to each other, clustering into two district groups representing the two species (Figure 1, Supplemental Table 1). While two genomes for DAOMC182701 were sequenced and assembled independently, one from each panel, the two genomes had >99.6% alignment, despite being sourced, cultured, sequenced, and assembled independently, supporting that the assemblies of both panels are of high quality (Figure 1). The percent genome alignment between, *P. pseudonodorum* and *P. nodorum* assemblies ranged from 58.0% to 70.0% (Figure 1), and were the best two aligning species within those pulled from NCBI, supporting that these species are closely related (Supplemental Table 1). Interestingly, some of the *P. pseudonodorum* isolates had reduced alignment to other *P. pseudonodoum* isolates from NCBI, indicating that there may be several different clades within the species or that some of these genomes may correspond to unique related species (Supplemental Table 1). The genomes of BC07, 07-SN-012, and 07-SN-017 were also somewhat larger than the other genomes (Table 1), which translated to reduced genome alignment (Figure 1); additional research will be needed to confirm that these cultures were pure and if they may contain elements that have been horizontally transferred between the species (Figure 1), with horizontal transfer being well-documented for *Parastagonaspora* spp. (McDonald et al., 2019). Another interesting observation was the low percent genome alignment of B102-44bF to other available genomes on NCBI (Supplemental Table 1). We extracted the ITS region from the genome and BLASTn analyses revealed that it may belong to the species *Phaeosphaeria pontiformis*, though additional analysis by morphology and additional barcoding regions would be needed to confirm the species assignment.

**Figure 1.**
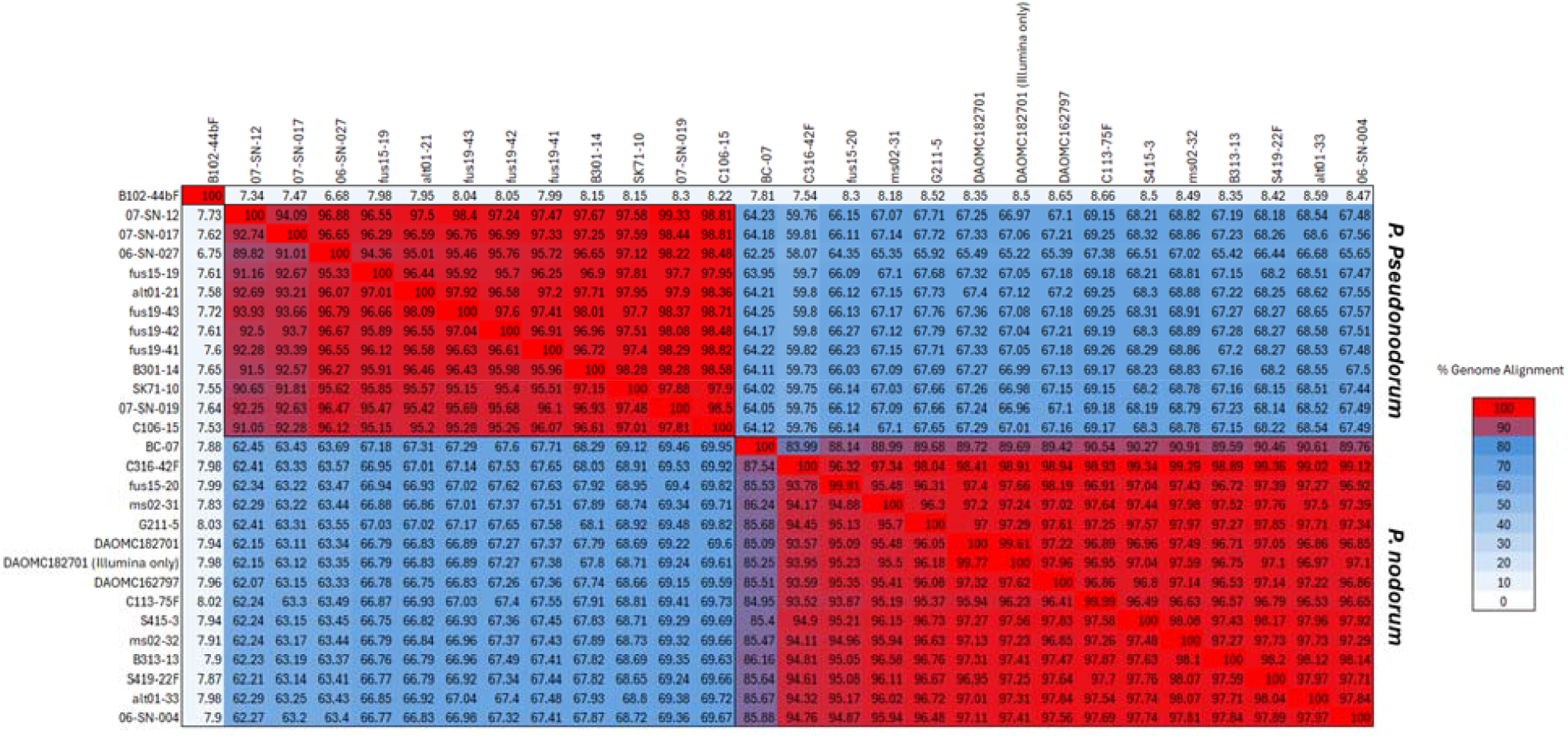
Percent genome alignment between the newly assembled genomes. Genomes were aligned using default parameters of nucmer and dnadiff. Percent genome alignment is indicated as a numerical colour gradient with white having no alignment, blue having intermediate alignment and red having high alignment. Data for alignment to additional Dothidiomycetes is in Supplemental Table 1.

Altogether, by making the genomes assemblies available here, and providing greater context to their source and identity, we are building upon a strong foundation of genomic resources that will support research on diseases that cause reduced yields in cereals in Canada and worldwide.

## Data Availability Statement

Data for the project are available in the NCBI databases under the BioProject accessions PRJNA1348192 and PRJNA1262417.

## Acknowledgements

We would like to acknowledge the DAOMC culture collection of Agriculture and Agri-Food Canada for providing isolates used in the study. Sequencing for the second panel was performed by the Centre d’expertise et de services Génome Québec (Montreal, Canada). Sequencing of the first panel was performed by the Canadian Grain Commission.

## References

Bertazzoni, S., Jones, D. A. B., Phan, H. T., Tan, K. C., & Hane, J. K. (2021). Chromosome-level genome assembly and manually-curated proteome of model necrotroph Parastagonospora nodorum Sn15 reveals a genome-wide trove of candidate effector homologs, and redundancy of virulence-related functions within an accessory chromosome. BMC Genomics, 22(1), 382. 10.1186/s12864-021-07699-8

Croll, D., Crous, P. W., Pereira, D., Mordecai, E. A., McDonald, B. A., & Brunner, P. C. (2021). Genome-scale phylogenies reveal relationships among Parastagonospora species infecting domesticated and wild grasses. Persoonia, 46, 116–128. 10.3767/persoonia.2021.46.04

Ficke, A., Cowger, C., Bergstrom, G., & Brodal, G. (2018). Understanding Yield Loss and Pathogen Biology to Improve Disease Management: Septoria Nodorum Blotch - A Case Study in Wheat. Plant Disease, 102(4), 696–707. 10.1094/PDIS-09-17-1375-FE

Gourlie, R., McDonald, M., Hafez, M., Ortega-Polo, R., Low, K. E., Abbott, D. W., Strelkov, S. E., Daayf, F., & Aboukhaddour, R. (2022). The pangenome of the wheat pathogen Pyrenophora tritici-repentis reveals novel transposons associated with necrotrophic effectors ToxA and ToxB. BMC Biology, 20(1), 239. 10.1186/s12915-022-01433-w

Griem-Krey, H., de Fraga Sant’Ana, J., Oggenfuss, U., Calegari-Alves, Y. P., Marques, A. L., Berger, M., Santi, L., Beys-da-Silva, W. O., & Habig, M. (2026). Transposable elements hitchhike on Starships across fungal genomes. Nature Communications, 17(1), 2634. 10.1038/s41467-026-69410-3

Hafez, M., Gourlie, R., Despins, T., Turkington, T. K., Friesen, T. L., & Aboukhaddour, R. (2020). Parastagonospora nodorum and Related Species in Western Canada: Genetic Variability and Effector Genes. Phytopathology®, 110(12), 1946–1958. doi:10.1094/PHYTO-05-20-0207-R

Kolmogorov, M., Yuan, J., Lin, Y., & Pevzner, P. A. (2019). Assembly of long, error-prone reads using repeat graphs. Nature Biotechnology, 37(5), 540–546. 10.1038/s41587-019-0072-8

Li, H. (2013). Aligning sequence reads, clone sequences and assembly contigs with BWA-MEM. arXiv preprint arXiv:1303.3997.

McDonald, M. C., Taranto, A. P., Hill, E., Schwessinger, B., Liu, Z., Simpfendorfer, S., Milgate, A., & Solomon, P. S. (2019). Transposon-Mediated Horizontal Transfer of the Host-Specific Virulence Protein ToxA between Three Fungal Wheat Pathogens. mBio, 10(5). 10.1128/mBio.01515-19

Robles-Zazueta, C., Crespo □ Herrera, L., Piñera □ Chavez, F., Rivera-Amado, C., & Aradottir, G. (2024). Climate change impacts on crop breeding: Targeting interacting biotic and abiotic stresses for wheat improvement. Plant Genome 17: e20365. In.

Seemann, T. (2026). Shovill: Robust Microbial Short-Read Genome Assembly. In Preprints: Preprints.

Thines, M., Crous, P. W., Aime, M. C., Aoki, T., Cai, L., Hyde, K. D., Miller, A. N., Zhang, N., & Stadler, M. (2018). Ten reasons why a sequence-based nomenclature is not useful for fungi anytime soon. IMA Fungus, 9(1), 177–183. 10.5598/imafungus.2018.09.01.11

Ueng, P., Bergstrom, G., Slay, R., Geiger, E., Shaner, G., & Scharen, A. (1992). Restriction fragment length polymorphisms in the wheat glume blotch fungus, Phaeosphaeria nodorum. Phytopathology, 82(11), 1302–1305. 10.1094/Phyto-82-1302

Ueng, P. P., & Chen, W. (1994). Genetic differentiation between Phaeosphaeria nodorum and P. avenaria using restriction fragment length polymorphisms. Phytopathology, 84(8), 800–806.

Ueng, P. P., Cunfer, B. M., Alano, A. S., Youmans, J. D., & Chen, W. (1995). Correlation between molecular and biological characters in identifying the wheat and barley biotypes of Stagonospora nodorum. Phytopathology, 85(1), 44–52.

Walker, B. J., Abeel, T., Shea, T., Priest, M., Abouelliel, A., Sakthikumar, S., Cuomo, C. A., Zeng, Q., Wortman, J., Young, S. K., & Earl, A. M. (2014). Pilon: An Integrated Tool for Comprehensive Microbial Variant Detection and Genome Assembly Improvement. PLOS ONE, 9(11), e112963. 10.1371/journal.pone.0112963

